# High-throughput direct screening of restriction endonuclease using microfluidic fluorescence-activated drop sorter based on SOS response in *E. coli*

**DOI:** 10.1101/2022.01.09.475563

**Authors:** Yizhe Zhang, Jeremy J. Agresti, Yu Zheng, David A. Weitz

## Abstract

A restriction endonuclease (RE) is an enzyme that can recognize a specific DNA sequence and cleave that DNA into fragments with double-stranded breaks. This sequence-specific cleaving ability and its ease of use have made REs commonly used tools in molecular biology since their first isolation and characterization in 1970s. While artificial REs still face many challenges in large-scale synthesis and precise activity control for practical use, searching for new REs in natural samples remains a viable route for expanding the RE pool for fundamental research and industrial applications. In this paper, we propose a new strategy to search for REs in an efficient fashion. Briefly, we construct a host bacterial cell to link the RE genotype to the phenotype of β-galactosidase expression based on the bacterial SOS response, and use a high-throughput microfluidic platform to isolate, detect and sort the REs. We employ this strategy to screen for the XbaI gene from constructed libraries of varied sizes. In single round of sorting, a 30-fold target enrichment was obtained within 1 h. The direct screening approach we propose shows potential for efficient search of desirable REs in natural samples compared to the conventional RE-screening method, and is amenable to being adapted to high-throughput screening of other genotoxic targets.

## INTRODUCTION

A restriction endonuclease (RE) is an enzyme that can recognize a specific DNA sequence and cleave that DNA into fragments with double-stranded breaks. This sequence-specific cleaving ability and its ease of use have made REs commonly used tools in molecular biology since their first isolation and characterization in the 1970s.^1–5^ With continuous improvement in flexibility and specificity,^6–14^ REs are gradually gaining traction as promising agents for targeted gene disruption in gene therapy against various virus infectious diseases, such as HIV and HPV.^15–18^ New REs with desirable features can be obtained through artificial synthesis or wild-type search. While artificial REs still face many challenges in large-scale synthesis and precise activity control for practical use,^8, 10, 19^ searching for new REs in natural samples remains a major route for expanding the RE pool to meet the growing needs in fundamental research and industrial applications.

The conventional RE-searching method utilizes modification methylases to indirectly screen for their companion REs. It predicts the potential RE genes by scanning through the sequence database for the companion modification methylase based on its conserved motif element, followed by the evaluation digestion tests to identify the new RE genes.^20, 21^ The major drawback of this indirect screening method is its dependence on the modification methylase, which precludes potential REs that do not have the companion methylases, for example, *Pac*I.^22^ Moreover, the scanning step requires the sample to be pre-sequenced, and the evaluation step requires intense labor, making it a challenge to search for REs in natural samples with potentially large library sizes.^23^ A few *in vitro* translation-based selection strategies have been proposed to directly search for new REs, but have had very limited searching range due to the stringent requirements on the recognition sequence.^24, 25^ Thus, new, more efficient methods to search for new REs would be valuable.

In this paper, we propose a new strategy to search for REs in an efficient fashion. We use a high-throughput screening strategy based on the bacterial SOS response:^26^ the presence of an RE gene is indicated by the over-expression of β-galactosidase (β-gal) through a specially designed host bacterial cell, which can be detected and sorted by a fluorescence-activated microfluidic drop sorter.^27^ We employ this strategy to screen for the XbaI gene from a constructed library and reach a 30-fold target enrichment for 0.1 % library within one hour.

## MATERIALS AND METHODS

### Host cell preparation

ER2745, an *Escherichia coli* (E. coli) derivative strain that contains a fusion of a DNA damage-induced SOS gene *dinD*^26^ and an indicator gene *lacZ*, is constructed as the host cell. The strain is deficient in all known endogenous restriction systems and expresses T7 RNA polymerase under *lac* control from a chromosomal location. To suppress the basal level expression of T7 RNA polymerase, ER2745 is transformed with pLysY.^28^ ER 2745/pLysY is then transformed with pTXB1_XbaI (*E. coli*^XbaI^ cell) or pTXB1_ΔXbaI (*E. coli*^ΔXbaI^ cell, with an internal deletion inside the XbaI gene) to compose the model library.

### Primer design

For easy identification, primers for the post-sorting PCR are designed to amplify a fragment of the insert such that the amplicons from the target and the control are short enough to run at a detectable distance on the gel without losing the base-pair difference information. Specifically, instead of amplifying the intact target insertion of 628 bp and the control insertion of 572 bp, the primers p_fwd_ (5’-TAGGGGAATTGTGAGCGGATAAC-3’) and p_rev_ (5’-GGAATCGGCCCTTGTTTTGATAG-3’) target for a 263-bp fragment in XbaI and a 207-bp fragment in ΔXbaI, so the shorter amplicons from the target and the control become well separated in the gel allowing for quantification.

### Cell culture

Host *E. coli* cells are cultured overnight in the standard LB medium with antibiotics Ampicillin (Amp) and Chloramphenicol (Cmp) (Sigma) until the OD_600_ reaches about 0.2. The harvested cells are spun down to remove the basal β-galactosidase (β-gal), and resuspended in LB medium and stored on ice for sample preparation.

### Sample preparation

For signal generation and detection in bulk, the harvested *E. coli*^XbaI^ cell and *E. coli*^ΔXbaI^ cell cultures are both induced with 0.5 mM (final concentration) of Isopropyl β-D-1-thiogalactopyranoside (IPTG; Invitrogen), and incubated at 37 °C in the dark for a given period of time before being disrupted, whereupon 1 mL of the cell suspension is spun down, resuspended in 0.7 mL sonication buffer (100 mM NaCl (Sigma), 25 mM Tris-HCl (Sigma), 1 mM β-mercaptoethanol (β-ME; Sigma); pH 8.0),^29^ then sonicated and spun down again to extract β-gal. 5 μL of the sonication extracts from *E. coli*^XbaI^ cell and *E. coli*^ΔXbaI^ cell are added to individual wells of a 96-well plate, where 50 μL of 0.2 mM Fluorescein-Di-β-D-Galactopyranoside (FDG; Life Technologies) is added as the fluorogenic substrate. The mixture is incubated at 37 °C in the dark for 20 min and then a fluorescence measurement is performed with a microplate reader (EM: 490 nm/AB: 514 nm; ThermoFisher Scientific).

For signal generation and detection in the droplet, a co-flow drop-maker microfluidic chip is used for better control over the onset of the enzymatic reactions.^30^ For the inner flow, the cells are mixed with Cmp (34 μg/mL), Amp (100 μg/mL) and Pluronic F127 (0.001%, to prevent cells adhering to the PDMS surface^31^) (Sigma) in LB to the density of 10^8^ cells/mL. For the middle flow, FDG is mixed in LB to 0.2 mM with IPTG (1 mM, final concentration), sodium *N*-lauroyl sarcosine solution (0.1%, cell lysate buffer to allow FDG in) (Sigma), Pluronic F127 (0.001%), Cmp (34 μg/mL) and Amp (100 μg/mL). The inner flow and the middle flow are infused at equal flow rates to form droplets that contain a mixture of LB with Cmp (34 μg/mL), Amp (100 μg/mL), Pluronic F127 (0.001%), FDG (0.1 mM), IPTG (0.5 mM) and sodium *N*-lauroyl sarcosine (0.05%). The average number of cells per drop is around 0.3, and the drop size is 23 μm in diameter.

### Microfluidic device fabrication

The microfluidic devices are fabricated by patterning channels in polydimethylsiloxane (PDMS) using conventional soft lithography methods.^32^ Briefly, for a 10-μm drop-maker in our experiments, SU8-3010 photoresist (MicroChem Corp.) is spin-coated onto the 3” silicon wafer and patterned by UV exposure through a photolithography mask. SU8-3025 photoresist is used for a 25-μm sorter. After baking and developing with SU-8 developer (propylene glycol methyl ether acetate; MicroChem Corp.), the 10-μm tall positive master of the drop-maker and the 25-μm tall positive master of the sorter are formed on the silicon wafers. Then a 10:1 (w/w) mixture of Sylgard 184 silicone elastomer and curing agent (Dow Corning Corp), degassed under vacuum, is poured onto the master and cured at 65 °C for 2 h. Afterwards, the structured PDMS replica is peeled from the master and inlet and outlet ports are punched in the PDMS with a 0.75-mm diameter biopsy punch (Harris Unicore). The PDMS replica is then washed with isopropanol, dried with pressurized air, and bonded to a 50 × 75 mm glass slide (VWR) through oxygen plasma treatment to form the device.

To fabricate the electrodes in the sorter, a 0.1-M solution of 3-mercaptotrimethoxysilane (Gelest) in acetonitrile (99.8%; Sigma) is flushed through the electrode channels and blown dry with pressurized air. A low-melting point solder (Indalloy 19 (52 In, 32.5 Bi, 16.5 Sn) 0.020” diameter wire; Indium Corp.) is infused into the electrode channels at 80 °C. An eight-pin terminal block with male pins (DigiKey) is then glued with Loctite 352 (Henkel) to the surface of the device. The solid electrodes in the shape of the channels are formed when the device is cooled down to room temperature. Electrical contacts are made with alligator clips and connected to a high-voltage amplifier (Trek) that is then connected to the function generator on an FPGA (field-programmable gate array) card (National Instruments).

To form the aqueous-in-oil emulsion, the microfluidic channels are rendered hydrophobic by flushing Aquapel (PPG Industries) through the channels and drying with pressurized air. To produce biocompatible stable drops, we dissolve 1.8% (w/w) EA surfactant (RainDance Technologies) in the fluorinated oil Novec HFE-7500 (3M).

### Microfluidic drop-making and sorting

To make the 23-μm-diameter drops, the cell suspension and substrate solution are infused into the inner and middle channels respectively of the 10 μm drop-maker at equal flow rates of 19 μL/h, while the oil is infused into the outer channel at 20 μL/h. The emulsion is collected in a 1-mL plastic syringe and incubated at 37 °C in the dark for 3 h to allow sufficient enzymatic reactions to take place, whereupon the drops are re-injected into a 25-μm sorter. The closely-packed drops are injected at 20 μL/h and are separated by the carrier oil, injected at 200 μL/h, producing drops which flow by the laser-situated detection window at a frequency of ~1.5 kHz, where their fluorescence intensity is interrogated using a photomultiplier tube and a custom LabView program. Drops with a fluorescence intensity above the defined threshold trigger the sorter to which we apply a single-ended electric square wave of 0.8~1.2 kV_pp_ using a frequency of 20 kHz, generated by the function generator on the FPGA. Approximately 5 cycles of the square wave are applied to deflect the selected drop into the collection channel using the dielectrophoretic force.^33^ The asymmetric design of the sorting junction defines the default flow to the waste channel when the electric field is not triggered.^27, 30^

### Colony PCR

Samples from the collection channel are washed in Perfluorooctane solution (20% (v/v) in HFE 7500; Sigma) to break the emulsion. Cells are retrieved in 50 μL nuclease-free water (Life Technologies). For a 25-μL PCR reaction, 2.5 μL of the cell suspension is mixed up with 1.25 μL of p_fwd_ (10 μM), 1.25 μL of p_rev_ (10 μM), 0.5 μL of dNTPs (10 mM; Life Technologies), 0.25 μL of Phusion Hot Start DNA polymerase (2 unit/μL; NEB), 5 μL of 5x Phusion buffer (NEB) and 14.25 μL of nuclease-free water. The inserted DNA fragments on the vector pTXB1 are fully amplified through a 60-cycle PCR process under the fine-tuned condition (initial denaturing: 98 °C, 3 min; denaturing: 98 °C, 10 s; annealing: 59.2 °C, 30 s; extension: 72 °C, 1 min).

### Gel electrophoresis

One μL of amplicons from colony PCR is mixed with 1 μL of Gel Loading Dye (6x; NEB) and 4 μL of nuclease-free water in each well on the 1.2% agarose gel (Sigma) stained with Ethidium Bromide (1 mg/μL; Life Technologies). 1 μL of 2-Log DNA Ladder (200 μg/mL; NEB) is mixed with 1 μL of Gel Loading Dye (6x) and 4 μL of nuclease-free water for the ladder well. The gel electrophoresis is running at 70 V in 0.5x TBE buffer (Life Technologies) containing 0.5 mg/μL Ethidium Bromide for 45 min.

## RESULTS AND DISCUSSION

### Restriction Enzyme Screening Strategy

We accomplished the direct RE screening in three steps: target isolation, signal generation, and target selection. Specifically, the host *Escherichia coli* (*E. coli*) cells that carry the library on the vectors are separated from each other into picoliter aqueous droplets after flowing through the microfluidic drop-maker from the inner channel; chemicals required by the enzymatic reactions for the signal-generation step such as Fluorescein-Di-β-D-Galactopyranoside (FDG) and Isopropyl β-D-1-thiogalactopyranoside (IPTG) are co-encapsulated with cells into the droplets through the middle channel on the co-flow device; fluorinated oil is flowed through the outer channel, as shown in Fig. 1a-1. This co-flow design of the device enables a better control over the onset of the enzymatic reactions. The collected emulsion is then incubated at 37 °C in the dark for a given period of time to allow the production of the fluorescence signal molecules through a series of enzymatic reactions in the target-carrying drops, as shown in Fig. 1a-2. The target-carrying fluorescent drops are then detected and sorted in the microfluidic sorting system, as shown in Fig. 1a-3.

**Fig.1.**
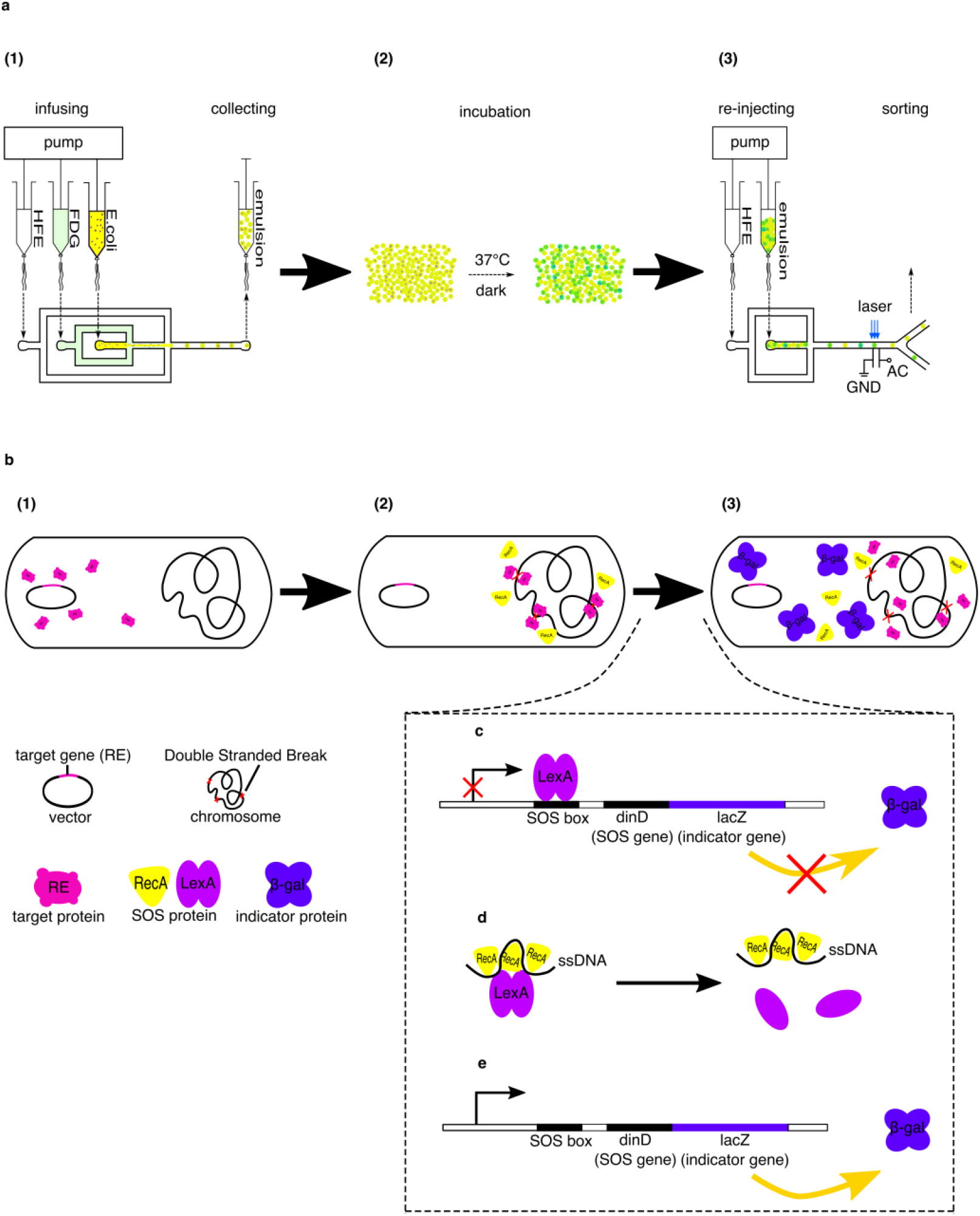
Restriction Enzyme (RE) screening strategy. **(a)** RE screening steps: (1) Co-encapsulation of the host cell *E. coli* and the fluorogenic substrate FDG. (2) Off-chip incubation for enzymatic reactions. (3) Detecting and sorting on the re-injected emulsion. **(b)** RE detection mechanism: Presence of RE in the host *E. coli* triggers the over-expression of β-galactosidase (β-gal) on the DNA-damage-induced pathway; β-gal can be fluorescently detected through its catalytic activity in the hydrolysis of the fluorogenic substrate. (1) RE expression. (2) Generation of DNA damage from RE activity induces SOS response that activates the up-regulation of RecA expression. (3) Over-expression of indicator protein β-gal. **(c-e)** SOS-regulated expression of indicator gene. For clarity, the symbols in the legend are not to scale.

To enable signal generation from the target drops, we utilize the host cell’s SOS response to activate the expression of the indicator protein. Specifically, we construct a host cell that lacks all known endogenous restriction systems and can express T7 RNA polymerase under *lac* control from a chromosomal location. We then fuse the indicator gene *lacZ* downstream to an SOS-inducible gene *dinD* (Fig. 1c). If the cell contains the target, REs are expressed upon the induction of IPTG during incubation, as illustrated in Fig. 1b-1. Functional REs cause DNA damage to the host cells due to their DNA cleavage activities, and thus trigger the cell’s SOS response in an attempt to repair the damage, leading to the over-expression of a myriad of SOS proteins,^26^ such as RecA (Fig. 1b-2). In the form of nucleoprotein filaments, RecA-ssDNA complex catalyzes the self-cleaving reaction of the repressor LexA (Fig. 1d), thus derepressing the *dinD :: lacZ* expression (Fig. 1c-1e) that leads to the production of the indicator protein β-gal (Fig. 1b-3). β-gal can be conveniently detected through the fluorogenic substrate FDG co-encapsulated in drops using our microfluidic fluorescence-activated drop sorter.^34^

### Signal-generation assay validation

To verify the SOS response-based signal-generation assay, we induce the enzymatic reactions in bulk and in drops respectively, and quantitatively examine the fluorescence signals through the detection experiments. We choose a typical RE XbaI (628 bp) as our target, and its truncated fragment ΔXbaI (572 bp) as the control.

For validation experiments in bulk, the target cells (*E. coli*^XbaI^) and the control cells (*E. coli*^ΔXbaI^) are both incubated for varied durations (0, 1, 2, 3, 4, 5 h) after IPTG induction, followed by the release of β-gal through sonication. The cell lysates from sonication are then added to the FDG-filled microtiter plate for fluorescence measurement. Without incubation, there is only a weak background fluorescence for both cell types; after incubation, *E. coli*^XbaI^ gives higher fluorescence intensity than *E. coli*^ΔXbaI^; as the incubation time is prolonged, the fluorescence intensity measured from *E. coli*^XbaI^ continues to increase whereas the measurement on *E. coli*^ΔXbaI^ shows no significant changes, as shown by the black and grey bars in Fig. 2a. The ratios of fluorescence intensity from *E. coli*^XbaI^ to that from *E. coli*^ΔXbaI^ at different incubation times, defined as the signal-to-noise ratio (s/n), are calculated from the measured data and plotted as the curve in Fig. 2a. The time-dependent strong fluorescence from *E. coli*^XbaI^ suggests the SOS-inducing DNA-cleavage activity of the intact XbaI, and hence validates the proposed SOS response-based signal-generation assay in bulk. We assume the background fluorescence from *E. coli*^ΔXbaI^ is probably due to the occasional relaxation of regulation on the *dinD::lacZ* pathway.

**Fig.2.**
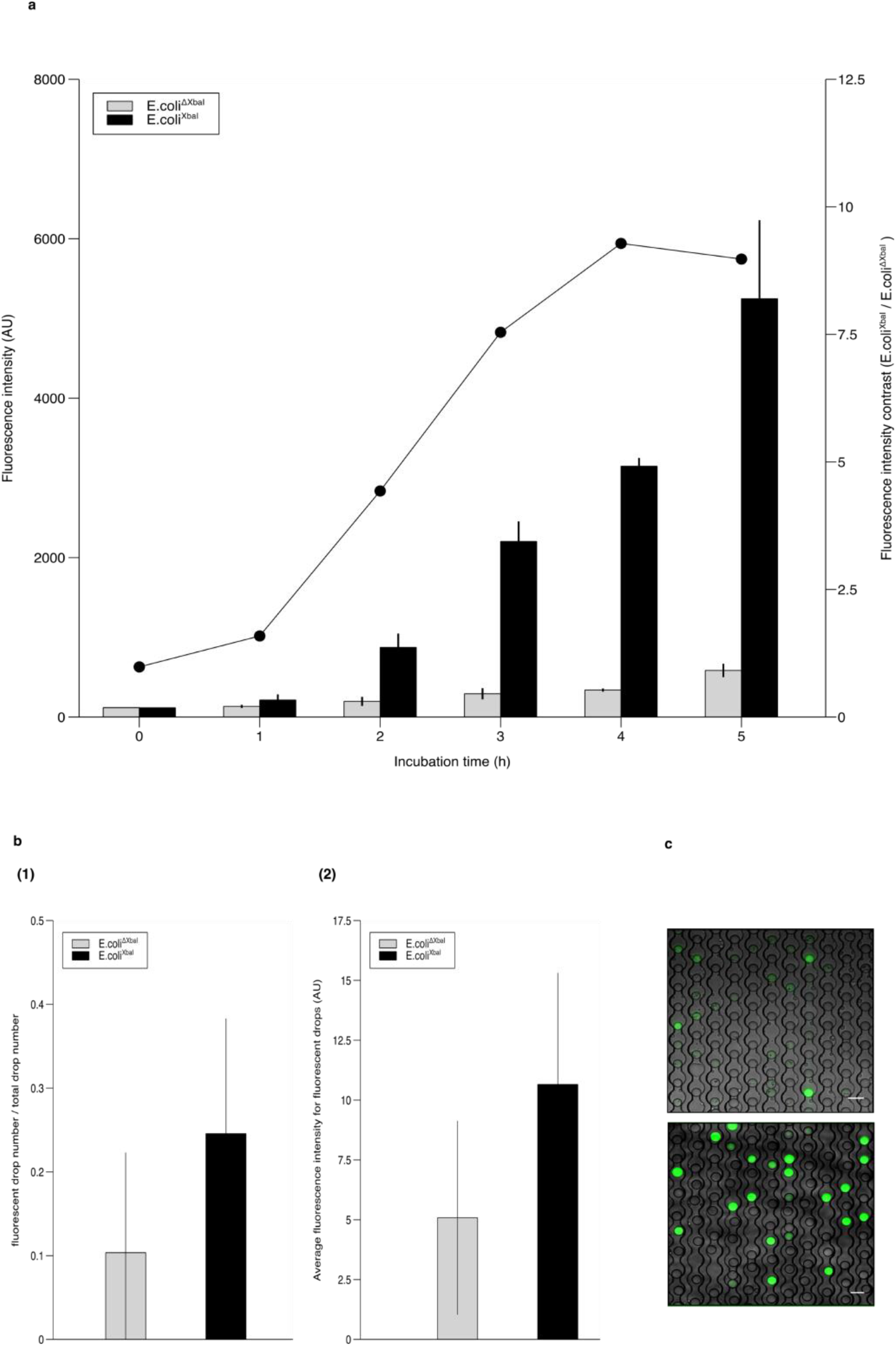
Signal-generation assay validation. **(a)** Fluorescence signal detection in bulk experiments. Bars: Fluorescence intensity from target cells (*E. coli*^XbaI^) and negative control cells (*E. coli*^ΔXbaI^) as a function of cell-incubation time. Curve: Fluorescence intensity contrast of *E. coli*^XbaI^ and *E. coli*^ΔXbaI^. **(b)** Fluorescence signal detection in drop experiments. Emulsion incubation time: 3 h. (1) Fraction of detectable fluorescent drops in total drops for *E. coli*^XbaI^ and *E. coli*^ΔXbaI^. (2) Average drop fluorescence intensity for *E. coli*^XbaI^ and *E. coli*^ΔXbaI^. **(c)** Representative drop fluorescence micrographs (overlaid with bright field images) in the signal detection experiments. The emulsion was allowed to flow into a “parking lot” device for imaging. Top: *E. coli*^ΔXbaI^ - encapsulated drops. Bottom: *E. coli*^XbaI^ - encapsulated drops. Scale bar: 40 μm. The average cell number per drop is kept at 0.3 in all droplet experiments.

For validation experiments in drops, we choose to generate drops with a diameter of 23 μm, a reliable size for single microbial isolation. To balance the ratio of single-cell drops to the number of empty drops, based on a Poisson distribution, we keep the ratio of the cell number to droplet number at 0.3 when preparing the cell suspension. To induce signal-generation in the drops, we incubate the emulsion at 37 °C for 3 h based on the results from the bulk detection experiments, at which point the signal is large enough (*s/n* = 7.5) for gating (Fig. 2a). Longer incubation raises the chance of drop coalescence without providing a significant improvement in the *s/n* level. One thing distinct from signal generation in bulk is the addition of sodium *N*-lauroyl sarcosine (sarkosyl) to the droplet to facilitate the cell’s uptake of FDG, since the mechanical rupture used in the bulk experiments is not used in the drops. To minimize the potential damage to cells, we carefully control the added sarkosyl to maintain a minimum concentration of 0.1 %.

As a quick test, we check the incubated drops under the fluorescence microscope and measure their fluorescence intensity from the fluorescence images using ImageJ (National Institutes of Health). For *E. coli*^XbaI^, the detectable fluorescent drops compose about 25% of the total drops (Fig. 2b-1, black), consistent with the Poisson distribution that predicts about 26% of the drops for a 0.3 *λ*-value. For *E. coli*^ΔXbaI^, as in the bulk experiments, we see some background fluorescence in the cell-carrying drops, but the fluorescence signal is weak in general: only 10% of the drops give detectable fluorescence (Fig. 2b-1, grey) and the average drop fluorescence intensity is less than 50% of that measured in the *E. coli*^XbaI^ experiments (Fig. 2b-2). The representative droplet images (overlay of bright-field and fluorescence images) from *E. coli*^ΔXbaI^ (Fig. 2c, top) and *E. coli*^XbaI^ experiments (Fig. 2c, bottom) are shown in Fig. 2c.

Both in bulk and in drops, we observe strong fluorescence signals from the target cells after the enzymatic activities in the cell are induced, which validates the proposed signal-generating assay on the basis of SOS response in the host cell.

### Sorting of XbaI

Upon validating the assay, we performed the sorting of the target RE, XbaI, from our model libraries. We construct our model libraries using *E. coli*^XbaI^ as target cells and *E. coli*^ΔXbaI^ as control cells. As a demonstration of the principle, four libraries with sizes of 1:2, 1:10, 1:100, and 1:1000 are prepared by mixing *E. coli*^XbaI^ and *E. coli*^ΔXbaI^ at ratios of 1:1, 1:9, 1:99, 1:999, respectively.

For each library, cells are encapsulated into aqueous drops along with IPTG, FDG and sarkosyl through the drop-making process. The collected emulsion is incubated off-chip in the dark at 37 °C for 3 h before being injected into the microfluidic sorter for detection and sorting. By monitoring the real-time distribution of the drop fluorescence intensity as calculated by LabView, we are able to instantly estimate the target-drop population and set an appropriate sorting threshold accordingly. Drops with fluorescence intensity above the threshold trigger the electric field and are deflected into the collection channel under the dielectrophoretic force.^33^ The consecutive snapshots from a fast-camera movie show the process of a target drop being deflected (indicated by white arrows in Fig. 3a). Compared to the non-target drop flowing through the sorting junction (Fig. 3a, the right drop in *t*_1_), the target drop (Fig. 3a, the right drop in *t*_3_) shows a distortion in shape and a shift in transversal position as a result of dielectrophoresis.

**Fig.3.**
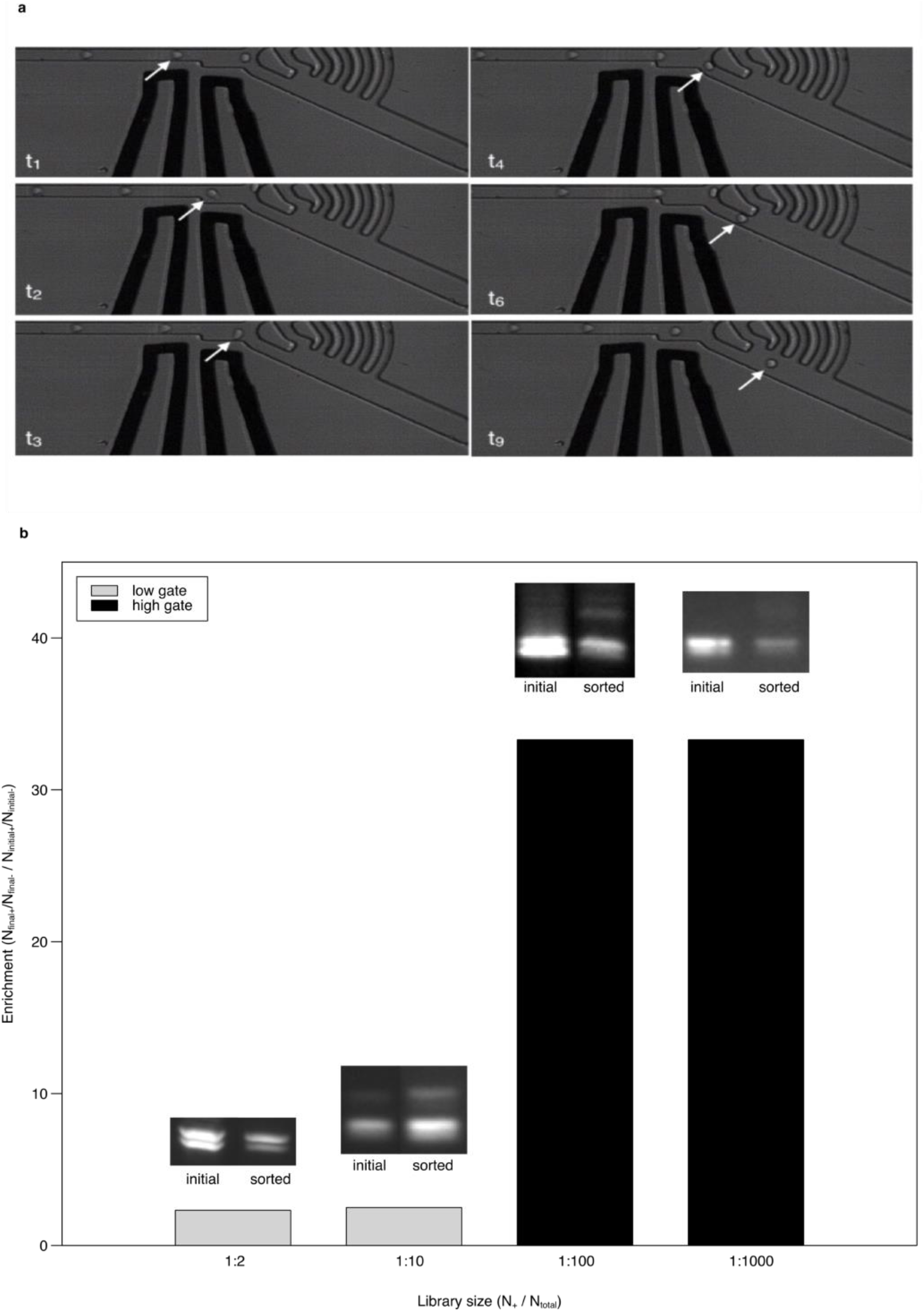
Microfluidic sorting of XbaI. **(a)** Snap shots of the sorting process on a target drop recorded by the fast camera (Photron Ultima512, 8000 fps). White arrows mark the target drop being deflected by dielectrophoresis. *t*_i_ (i = 1, 2, 3, 4, 6, 9) indicates the time points when the snap shots were taken. *t*_i+1_ - *t*_i_ = 0.125 ms. **(b)** The single-round sorting enrichment obtained for varied libraries. Insets: agarose gel pictures of the PCR amplicons from the initial and sorted samples. Software: ImageJ (National Institutes of Health).

We analyze the sorting result by evaluating the target enrichment in the sorted sample using gel electrophoresis. Briefly, the inserted DNA fragments XbaI or ΔXbaI on the vector pTXB1 from the sorted sample are recovered into distilled water after oil removal, and then amplified by colony PCR. The amplicons from the two templates are then separated by length on the agarose gel through electrophoresis. By quantifying the band brightness using ImageJ, we can obtain the ratio of the concentrations of XbaI and ΔXbaI in the sorted sample. We define our sorting enrichment as:

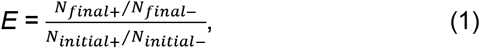

where *E* is the enrichment, *N*_final+_ is the number of target DNA insertion (XbaI) in the sorted sample, *N*_final-_ is the number of truncated insertion (ΔXbaI) in the sorted sample, *N*_initial+_ and *N*_initial-_ represent the numbers of XbaI and ΔXbaI in the unsorted sample.

The enrichment from a single round of sorting for libraries of 1:2 and 1:10 is about 2.5 fold at a low sorting threshold, as shown by the gray bars in Fig. 3b. When the sorting threshold is raised for libraries of 1:100 and 1:1000, the enrichment is increased to around 33.3, as shown by the black bars in Fig. 3b. Agarose gel pictures from unsorted and sorted samples for each library are shown as the insets in Fig. 3b, where the top band in each lane is from XbaI and the bottom band corresponds to ΔXbaI.

Ideally, if the fluorescence is only generated from the target, the enrichment would be determined by the distribution of the targets over the drops upon encapsulation, which, in our drop experiments, is the Poisson distribution. Therefore, merely by diluting the cell suspension to minimize the drops with more than one cell, the enrichment can be improved.^27^ In fact, we observe in our experiments that control cells also generate a background fluorescence, which we assume is due to some leakiness of the *dinD::lacZ* pathway, so a higher sorting threshold can also effectively suppress the number of the false positive drops and lead to a higher enrichment, as demonstrated in Fig. 3b. However, improving the enrichment by diluting the cell suspension or increasing the sorting threshold requests longer collection time to obtain sufficient samples for post-sorting processing. Thus, the high sorting threshold we chose in our experiments is a trade-off between enrichment and efficiency, which allows us to reach a reasonable enrichment within an hour of sorting for our model libraries of up to 1:1000 in size. For screening of larger libraries, it would be more practical to run a multi-round sorting than to optimize the enrichment with single-round sorting through modulating the sorting threshold or the cell density.

## CONCLUSIONS

We demonstrate the principle of an SOS response-based direct RE-screening strategy on the microfluidic platform through successful enrichment of the XbaI gene from model libraries with varied sizes. Compared to the conventional RE-screening method,^21^ our direct screening approach shows the potential for efficient search of desirable REs in natural samples. It does not rely on the modification methylase and therefore has a broader searching range; it does not involve any database-scanning and thus not requires pre-sequencing for the sample; and it runs on a microfluidic platform and therefore is highly cost- and time-efficient.

## ACKNOWLEDGEMENTS

This work was supported by the National Science Foundation (DMR-1310266 to D.A.W.) and the Harvard Materials Research Science and Engineering Center (DMR-1420570 to D.A.W.). We thank Dr. Richard J. Roberts at New England BioLabs for critical reading of the manuscript.

